# GeNNet: An Integrated Platform for Unifying Scientific Workflow Management and Graph Databases for Transcriptome Data Analysis

**DOI:** 10.1101/095257

**Authors:** Raquel L. Costa, Luiz M. R. Gadelha, Marcelo Ribeiro-Alves, Fabio Porto

## Abstract

**Background:** There are many steps in analyzing transcriptome data, from the acquisition of raw data to the selection of a subset of representative genes that explain a scientific hypothesis. The data produced may additionally be integrated with other biological databases, such as Protein-Protein Interactions and annotations. However, the results of these analyses remain fragmented, imposing difficulties, either for posterior inspection of results, or for meta-analysis by the incorporation of new related data. Integrating databases and tools into scientific workflows, orchestrating their execution, and managing the resulting data and its respective metadata are challenging tasks. Running in-silico experiments to structure and compose the information as needed for analysis is a daunting task. Different programs may need to be applied and different files are produced during the experiment cycle. In this context,the availability of a platform supporting experiment execution is paramount.

**Results:** We present GeNNet, an integrated transcriptome analysis platform that unifies scientific workflows with graph databases for selecting relevant genes according to the evaluated biological systems. GeNNet includes pre-loaded biological data, pre-processes raw microarray data and conducts a series of analyses including normalization, differential expression inference, clusterization and geneset enrichment analysis. To demonstrate the features of GeNNet, we performed case studies with data retrieved from GEO, particularly using a single-factor experiment. As a result, we obtained differentially expressed genes for which biological functions were analyzed. The results are integrated into GeNNet-DB, a database about genes, clusters, experiments and their properties and relationships.The resulting graph database is explored with queries that demonstrate the expressiveness of this data model for reasoning about gene regulatory networks.

**Conclusions:** GeNNet is the first platform to integrate the analytical process of transcriptome data with graph database. It provides a comprehensive set of tools that would otherwise be challenging for non-expert users to install and use. Developers as well can add new functionality to each component of GeNNet. The resulting data allows for testing previous hypotheses about an experiment as well as exploring new ones through the interactive graph database environment. It enables the analysis of different data on humans, rhesus, mice and rat coming from Affymetrix platforms.

## Background

The growing accumulation of molecular biology data motivated the development of pipelines, workflows and platforms for analyzing data. Many researchers are using these integrative approaches for analyzing metagenomes, proteomes, transcriptomes and other ‘omics’ data. For transcriptomes, microarray and RNA-seq are currently the main technologies available and widely used. The low cost of microarray, in relation to RNA-seq, still makes its use very appealing for well-known organisms. Regardless of the technology, there are many steps from the acquisition of raw data to the selection of a subset of representative genes that explain the hypothesis of the scientists. Furthermore, these genes can be grouped based on their gene expression pattern, to which biological function can attributed. The results of gene expression analysis may bring new insights to the discovery of new targets for drug development as well as for uncovering novel biological functions and mechanisms.

However, the results of these analyses remain fragmented, imposing difficulties, either for posterior inspection of results, or for meta-analysis by the incorporation of new related data. Integrating databases and tools into computational analyses, orchestrating their execution, and managing the resulting data and its respective metadata are challenging tasks [12]. Academic journals are demanding better reproducibility of computational research, requiring a precise record of parameters, data and processes (also called provenance [5]) used in these activities to support validation by peers [27].

Overcoming many of these challenges can be supported by designing and executing these computational analyses as scientific workflows [6], which consist of compositions of different scientific applications. Their execution is usually chained through data exchange, i.e. data produced by an application is consumed by subsequent applications. Scientific workflow management systems (SWMSs) enable for managing the life cycle of scientific workflows, which is usually given by composition, execution and analysis [21]. Many SWMSs, such as Taverna [23] and Swift [32], natively support gathering provenance [10] and executing scientific applications on scalable computational resources [31] such as high performance computational clusters and cloud computing infrastructures.

The heterogeneity of biological data makes its representation with a conceptual data schema that follows a fixed and strict structure, such as in relational databases challenging. Modifying the data schema in these cases can result in conflicts or inconsistencies in a database. In the era of expanding and interconnected information, new data models appeared such as column-oriented, key-value, multidimensional, and graph databases. These are commonly called NoSQL (*Not only SQL*) [30] databases and often have advantages in terms of scalability. Graph-based data models, in particular, are useful for data in which the relationship between attributes is one of the main aspects to be taken into consideration during querying. The graph database is an intuitive way for connecting and visualizing relationships. In graph databases the nodes represent objects and the edges represent the relationships between them. Both, nodes and edges can hold properties, which add information about the objects or the relationships. We chose the graph data model since it is the most adequate to represent the results in a natural way focusing on interactions. In recent years, this database model has been used in many different bioinformatics applications and are particularly promising for biological data sets [25, 17, 3, 14, 22]. Have and Jensen [13] observed that for path and neighborhood queries, Neo4j, a graph database, can be orders of magnitude faster than PostgreSQL, a widely used relational database, while allowing for queries to be more intuitively expressed.

Integrating scientific workflows with database systems becomes a powerful framework, in which scientists can express complex data pre-processing analysis and make available for further investigation treated data to be queried using a high-level query language. We argue that integrated web applications, involving scientific workflows and databases, can hide the complexity of underlying scientific software by abstracting away cumbersome aspects, such as managing files and setting command-line parameters, leading to increased productivity for scientists. One important aspect of enabling reproducible computational analyses is keeping track of the computational environment components, i.e., operating system, libraries, software packages and their respective versions. Operating system-level virtualization through the use of software containers allows for creating within an operating system instance isolated environments that behave like a server. These isolated environments, also called containers, can be built in a programmable way to ensure that they will be composed by the same libraries and software packages every time they are instantiated.

In this paper, we present GeNNet, an integrated transcriptome analysis platform that unifies scientific workflows with graph databases for determining genes relevant to evaluated biological systems. GeNNet includes pre-loaded back-end data, pre-processes raw microarray data and conducts a series of analyses including normalization, differential expression, annotation, clusterization and functional annotation. During these analyses, the results are stored in different formats, e.g., figures, tables and R workspace images. Furthermore, the results are stored as a graph database that can be persisted for the user. The graph database represents networks that can be explored either graphically or using a flexible query language. The application additionally offers an easy-to-use web interface tool developed in Shiny^1^ for automated analysis of gene expression. The implementation follows best practices for scientific software development [33], for instance, by recording provenance information and using software containers to distribute the platform and allowing for portability and reproducibility. As far as we know, GeNNet is the first platform for transcriptome data analysis that tightly couples a scientific workflow with a persistent biological (graph) database while supporting reproducibility through the use of provenance

## Implementation

GeNNet innovates in its use of a graph-structured conceptual data model coupled with scientific workflow management, software containers for portability and reproducibility, and a productive and user-friendly web-based front-end (see in *framework*). In the following subsections we describe these components in detail: workflow (GeNNet-Wf), graph database (GeNNet-DB), and web application (GeNNet-Web).

**Figure 1:**
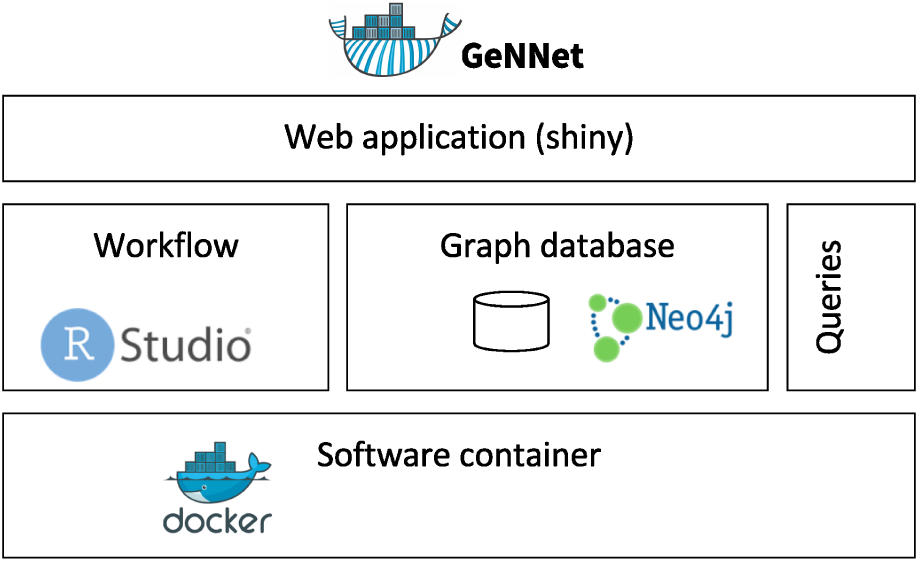
GeNNet framework with the components as GeNNet-Wf, GeNNet-DB and GeNNet-Web.

### Workflow

GeNNet-Wf was modularized in two main stages: background preparation and execution of workflow activities (Figure 2). The *background preparation* stage is executed during the construction of the GeNNet software container (described in section *container*), the resulting data is ready for use when the GeNNet platform is started. The *workflow activities* stage is comprised of the execution of a series of tools and libraries to analyze the transcriptome data uploaded by the user in conjunction with the background data.

**Figure 2:**
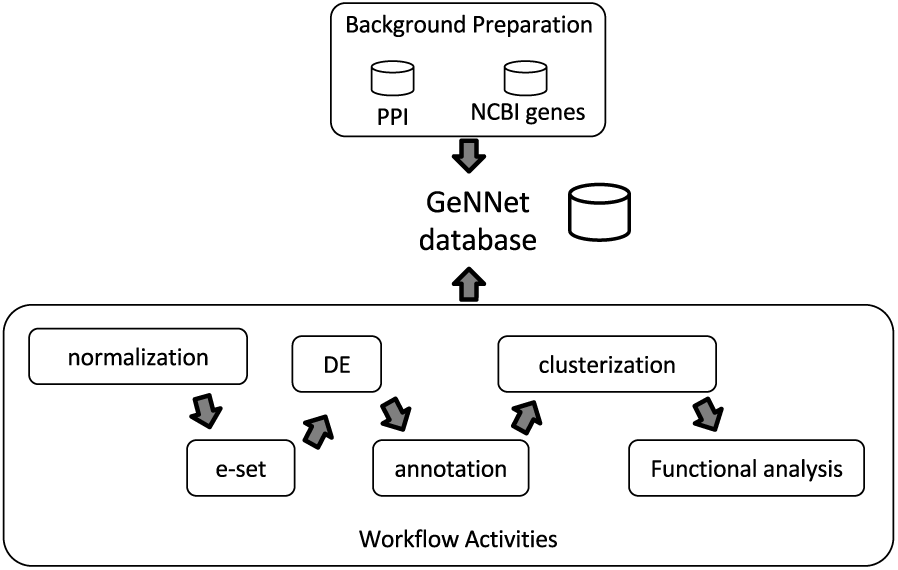
Workflow scheme represented by two stages, *Background Preparation* in the top and *Workflow Activities* in the bottom. Results of both stages are loaded to the GeNNet database. The *Workflow Activities* stage is shown with its different steps of the analysis.

### ‘Background’ Preparation

The genes and existing relationships among them along with other associated elements are the core of transcriptome analysis. In our platform, we build a data layer called *background* that contains information about all the genes annotated/described for the organisms targeted by GeNNet. Therefore, the background constitutes an independent and self-contained layer of the experiment. The independence of this information, generated during the construction of the software container, enables gains in efficiency in populating the GeNNet database since the data is bulk-inserted. In this version of the platform, the background data is comprised of two main sources: (i) gene information about human, rhesus, mice and rat, obtained from NCBI annotations [28] and (ii) Protein-Protein Interaction (PPI) network, retrieved from STRING-DB (Search Tools for Retrieval of Interacting Gene/Proteins) [9] (version 10). All genes imported from NCBI become nodes if the graph database and some of the main information associated to them (such as symbol, entrezId, description, etc.) are modeled as node properties. The information derived from STRING-DB PPI become edges (relationships between genes) with score values associated to the nodes (neighborhood, gene fusion, co-occurrence, co-expression, experiments, databases and text-mining). This layer of data is added to the graph database during the construction of the GeNNet container; more detail about the representation and implementation can be found in section *graphdb.*

### Workflow Activities

The *workflow activities* step (Figure 2) consists of a series of steps executed sequentially. This module was written in R using different packages mainly from the Bioconductor [7] and CRAN repositories. The steps are detailed next.

#### Normalization

This step consists in normalizing the raw data from an informed Affymetrix platform using either RMA [16] or MAS5 methods, both available in the affy [11] package. During this step, some quality indicator plots are generated (as boxplot of probe level, Spearman correlation and density estimates) as well as a normalized matrix (log-normalized expression values).

#### e-set

In this step, data about the experimental design should be added along with log-normalized expression values. This generates an ExpressionSet (eSet) object, a data structure object of the S4 class used as base in many packages developed in Bioconductor transcriptome analysis. This format gives flexibility and access to existing functionality. The input file must be structured using mainly two columns: a column named SETS for the experimental design, and a column named SAMPLE_NAME for the names of the files containing raw sample expression matrix data.

#### Filtering/Differential expression inference

Differential expression (DE) inference analysis allows for the recognition of groups of genes modulated (up- or down-regulated) in a biological system when compared against one or more experimental conditions. In many situations this is a core step of the analysis and there are a great diversity of experimental designs (such as control versus treatment, consecutive time points, etc) allowing the inference. In our platform, we use the limma package to select the DE genes [29] on single-factor experimental designs based on a gene-based hypothesis testing statistic followed by a correction of multiple testing given by the False Discovery Rate (FDR) [18]. Furthermore, a subset of DE genes can be selected based on a up- and down-regulation, expressed as a logarithmic (base 2) fold-change (logFC) threshold. Results of this step are displayed as Volcano plots and Matrices containing the DE genes.

#### Annotation

The annotation step consists of annotating the probes for the corresponding genes according to the Affymetrix platform used in the experiment.

#### Clusterization

This step consist in analyzing which aggregated genes have a similar pattern (or level) of expression. We incorporated clusterization analysis including hierarchical methods, *k*-medoids from the package PAM (Partitioning Around Medoids) [26] and WGCNA (Weighted Gene Coexpression Network Analysis) [19].

#### Functional Analysis

Genes grouped by similar patterns enables the identification of over-represented (enriched) biological processes (BP). In our approach we conducted enrichment analyses applying hypergeometric tests (with p-value < 0.001) as implemented in the GOStats package [8]. The universe is defined as the set of all genes represented in a specific Affymetrix platform, or, in case of multiple platforms in a single experiment design, the universe is defined as the common and unique genes in among all Affymetrix platforms. The subset, geneset, is defined either by the set of diferentially expressed (DE) genes between a test and a control condition (control versus treatment design), or by the union of the DE genes selected among the pairwise comparisons among groups in all other single-factor experimental designs. Ontology information for the gene and universe sets is extracted from the Gene Ontology Consortium database [2].

#### Execution

GeNNet is designed to automatically execute the workflow through the web application interface (available at http://localhost:3838/gennet, when the software container is running). However, users that intend to implement new functions or even execute the workflow partially, can use the RStudio server interface in GeNNet (through at http://localhost:8787 after starting the software container). More details are available in Supplementary Material.

### Graph database

GeNNet database (GeNNet-DB) schema is based on the Neo4j database management system, a free, friendly-to-use and with broad community support graph database, with its nodes, edges and relationships. Although a NoSQL database has no fixed schema, we defined an initial schema to help and guide the GeNNet-DB (Figure 3). Vertices and edges were grouped into classes, according to the nature of the objects. We defined the labels as GENE, BP (Biological Process), CLUSTER, EXPERIMENT, ORGANISM, and a series of edges as illustrated in Figure 3. In the GeNNet platform there is an initial database defined by interactions between genes as described in *Background preparation.* During the execution of GeNNet-Wf, using Shiny or RStudio, new nodes and connections are formed and added to the database. The resulting information is stored in the graph database using the RNeo4j package^2^. It can also be accessed directly through the Neo4j interface (available at: http://localhost:7474). It is possible to query and access the database in this interface using the Cypher language, a declarative query language for Neo4j, or Gremlin, a general-purpose query language for graph databases. These query languages allow for manipulating data by updating or deleting nodes, edges and properties in the graph. Querying also allows for exploring new hypotheses and to integrate new information from different resources that are related to the targeted experiment. GeNNet-DB is persistent and the resulting database is exported to a mounted directory. Its contents can be loaded to a similar Neo4j installation. For further details one can read the Neo4j manual.

**Figure 3:**
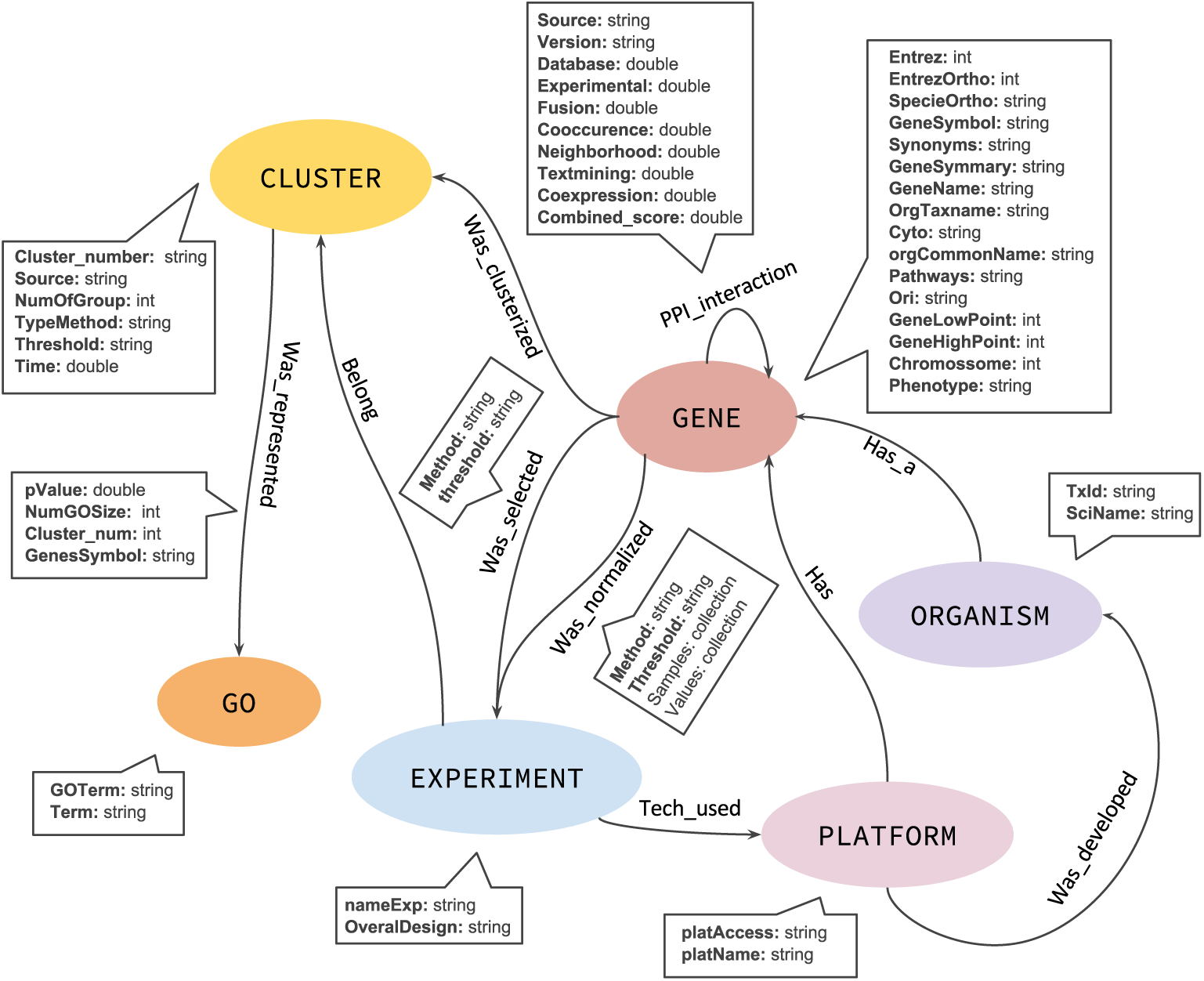
The graph database schema representing the nodes (in oval), relationships (arrows). The descriptive boxes showing the mainly properties in nodes and edges.

### Web application

GeNNet-Web provides a user-friendly way to execute GeNNet-Wf. We developed an easy-to-use layout for providing the parameters and automatically executing all steps of the workflow experiment. The parameters comprise the input of the web application, which include: descriptors for experiment name and overall design; type of normalization; differential expression settings; experiment platform and organism; and clusterization method. After executing GeNNet-Wf, GeNNet-Web allows for easy retrieval and visualization of its outputs, which are given by a heatmap, graph database metrics (e.g., number of nodes, number of edges, relationships between nodes), and the list of differentially expressed genes selected. In addition to the outputs generated in the web application, the underlying workflow generates the output files as described in subsection *gennet-workflow*.

### Software container

GeNNet was built on top of the Docker^3^ software containerization platform. This enables users to download a single software container that includes all the components of GeNNet and behave the same way independently of the hosting operating system. The software container was successfully tested on CentOS Linux 7, Ubuntu Linux 14.04, MacOS X 10.11.6 and Windows 10. The software container for GeNNet, specified in a script named *Dockerfile,* was built according to the following steps: (i) The operating system environment is based on CentOS Linux 7 with software packages required by GeNNet, such as R (v. 3.3.1), installed from the official CentOS repository and the EPEL (*Extra Packages for Enterprise Linux*) repository; (ii) The R packages required by GeNNet, installed from the CRAN repository; (iii) RStudio (v. 1.0.44) server and the Neo4j (Community Edition v.3.0.6) graph database, installed from their respective official repositories; (iv) Supporting data sets, such as PPI, loaded to the graph database; (v) GeNNet-Wf, implemented in R, installed in RStudio; (vi) Shiny, a web application server for R, installed from its official repository. GeNNet-Web, which calls GeNNet-Wf, is loaded to Shiny.

**Figure 4:**
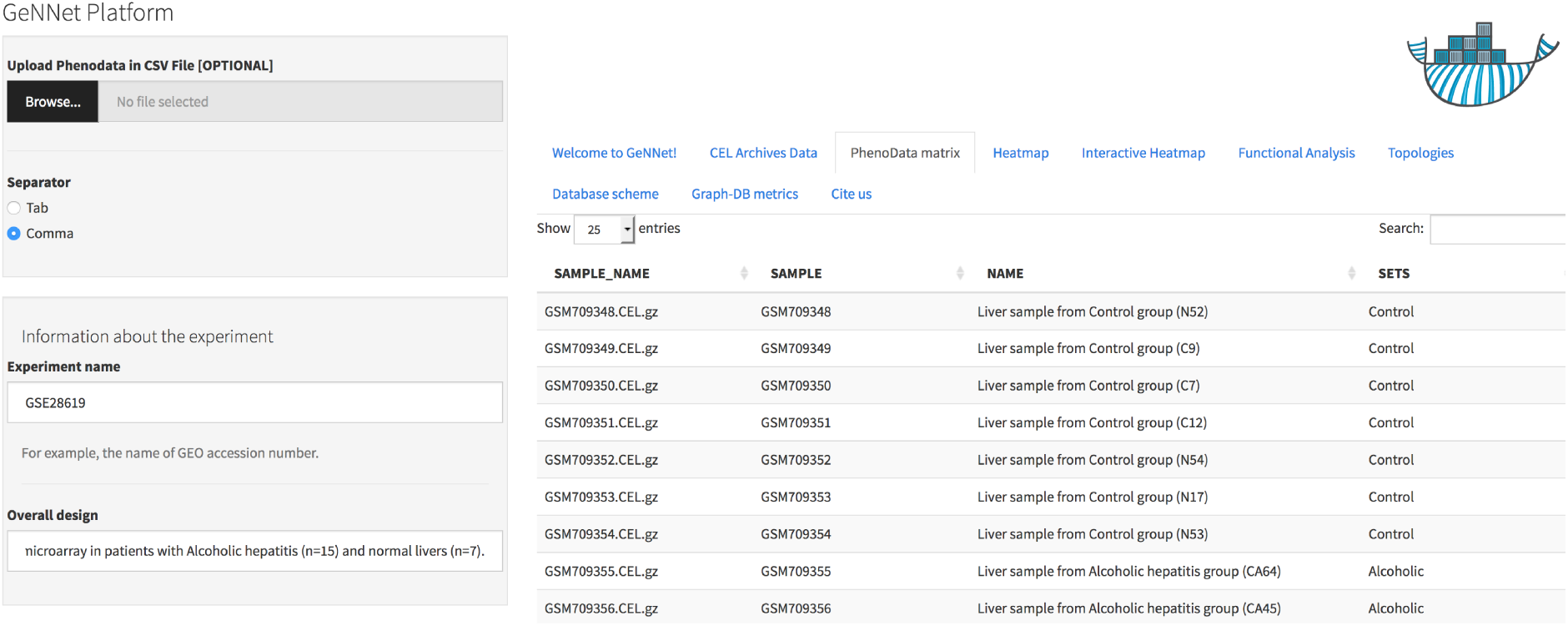
User-friendly interface in GeNNet. Left hand side showing parameters settings and right hand side showing some tables, figures and results.

### Computational experiment reproducibility

Reproducibility is accounted in GeNNet in two aspects. Firstly, the platform provides a provenance trace record generated by the RDataTracker package [20] for R. The trace contains the activities executed by the workflow and the data sets consumed and produced by them. This trace is exported to a persistent directory. Secondly, the adoption of software containers allows for using the same environment (operating system environment, libraries, and packages) every time GeNNet is instantiated and used. Both the provenance trace and the preservation of the execution environment with software containers significantly help the computational experiment reproducibility since users can retrieve from the former the parameters and data sets used in analyses and, from the latter, re-execute them in the same environment, as provided by the GeNNet software container.

## Results

### Experimental data — Use case scenarios

To demonstrate our application, we selected some case studies to be analyzed on GeNNet. The data was obtained from GEO [4] and agreed with the following criteria: (i) raw data availability; (ii) microarray data coming from the Affymetrix platform; (iii) encompassing humans, rhesus, mice and rat organisms; (iv) having single-factor experiment design. The datasets retrieved for validating our platform are listed in Additional material 3. Further details about each experiments can be found in the original articles.

As an example of a specific and more detailed case study, we re-analyzed a gene expression experiment from a patient with alcoholic hepatitis (15 samples in total) versus healthy individuals (7 samples in total) [1]s. The data for this experiment was obtained from GEO with accession number GSE28619. The study used the Affymetrix Human Genome U133 Plus. Data was normalized using the MAS5 method and the differential expressed gene selection criteria were FDR < 0.05 and absolute log2(Fold-Change) > 1. The genes were clustered using the Pearson correlation method as measure of dissimilarity. Next, the clusters were associated to biological functions through the hypergeometric test (with p-value < 0.001 as threshold). As a result, 2.478 differentially expressed genes were obtained and 513 ontological terms were represented (p-value < 0.001). A major part of the analytical process resulting information was incorporated to GeNNet-DB and beside the database, the results were exported to different formats such as figures (heatmaps, boxplots, etc.), tables and provenance (Figure 5).

**Figure 5:**
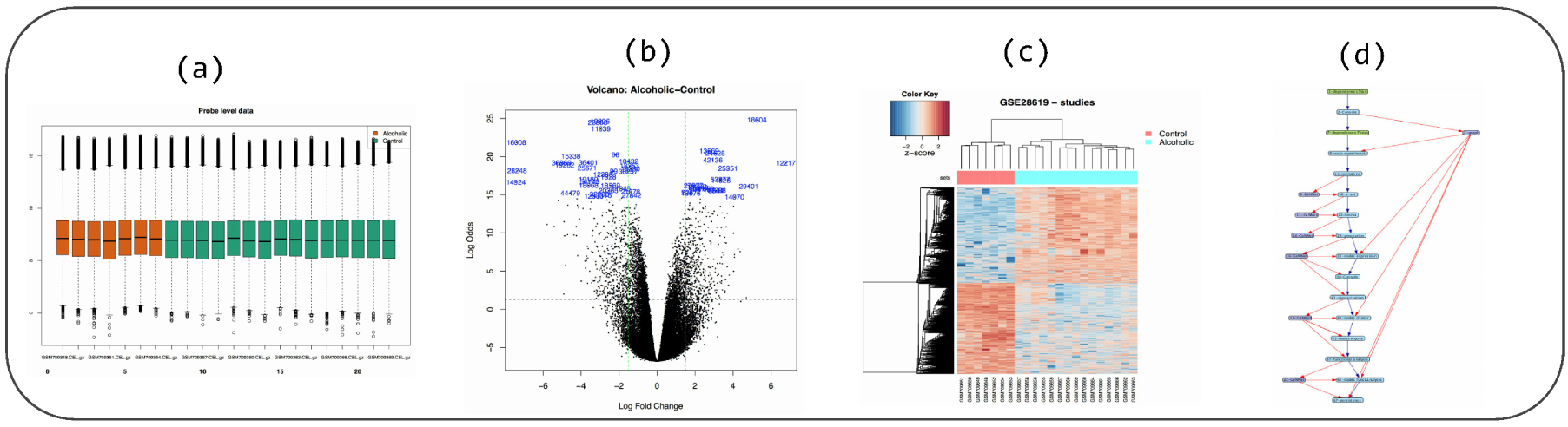
Some figures generated during workflow execution: (a) boxplot (quality indicator), (b) volcano plot and (c) heatmap. In (d), the provenance trace of a GeNNet-Wf execution is represented as a data derivation graph (DDG).

The database generated during GeNNet-Wf execution facilitates data representation as interaction networks, in an approach that allows for exploring a great variety of relationships among its composing entities, besides making new insights for subnetwork exploration possible. Depending on the type of these interactions, different kinds of networks and topologies can be defined and analyzed. Through the data representation used in GeNNet-DB traversal queries are possible. We illustrate a typical scenario for which the user just needs to query GeNNet-DB to solve them. Using the Cypher declarative query language with direct access to the database, we formulated some demonstration queries using as example the dataset analyzed above.

**Query 1:**
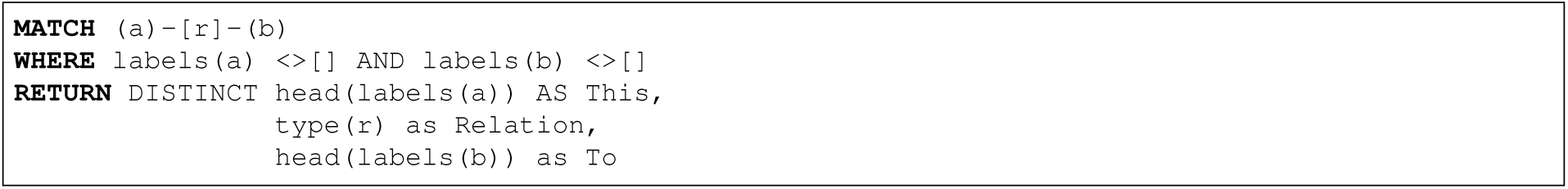
What are the existing relationships among nodes in the database? This is a simple query that returns all existing relationships among different node labels and types. The result of the query was represented as a graph in Figure 6.

**Figure 6:**
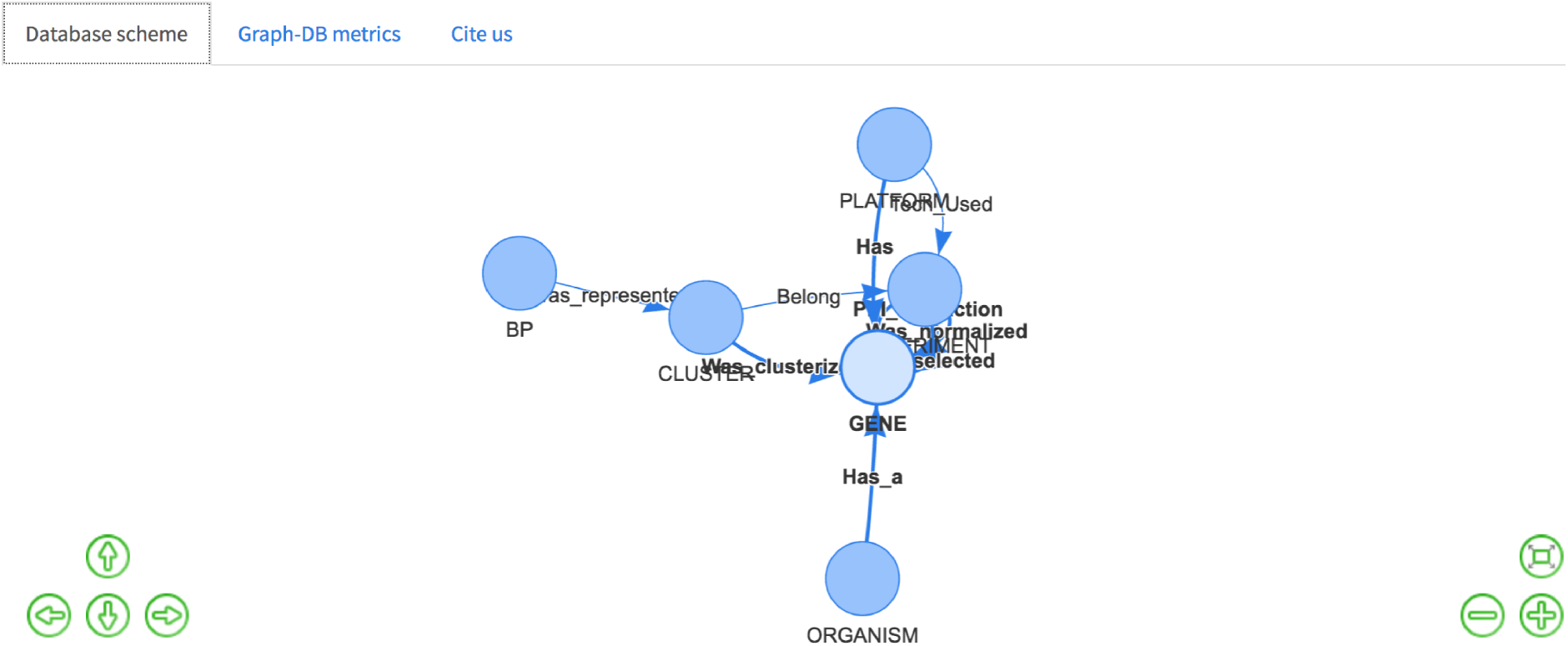
Database schema with all the existing nodes and relationships.

**Query 2:**
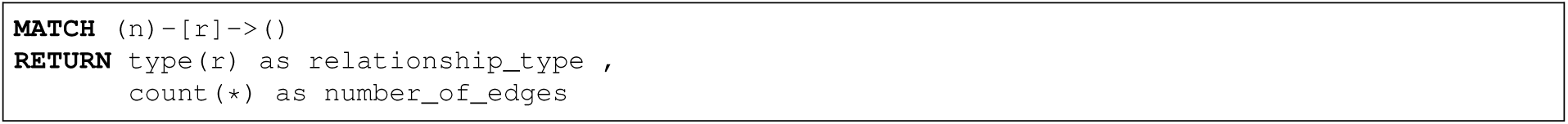
What is the number of edges per type of relationship? This query returns the number of edges according to each type of edge (1). In the result, one can observe the number of normalized and annotated genes (Was_normalized = 21031) as well as the nuber of DE genes found using the methodology selected (Was_selected = 2478) for the dataset analyzed.

**Table 1:** Types of edges with the respective number of edges stored in GeNNet-DB.

| relationship_type | number_of_edges |
| --- | --- |
| Was_represented | 513 |
| Was_clusterized | 10116 |
| PPI_interaction | 3471455 |
| Was_selected | 2478 |
| Has_a | 194588 |
| Belong | 2 |
| Tech_Used | 1 |
| Was_normalized | 21031 |

**Query 3:**
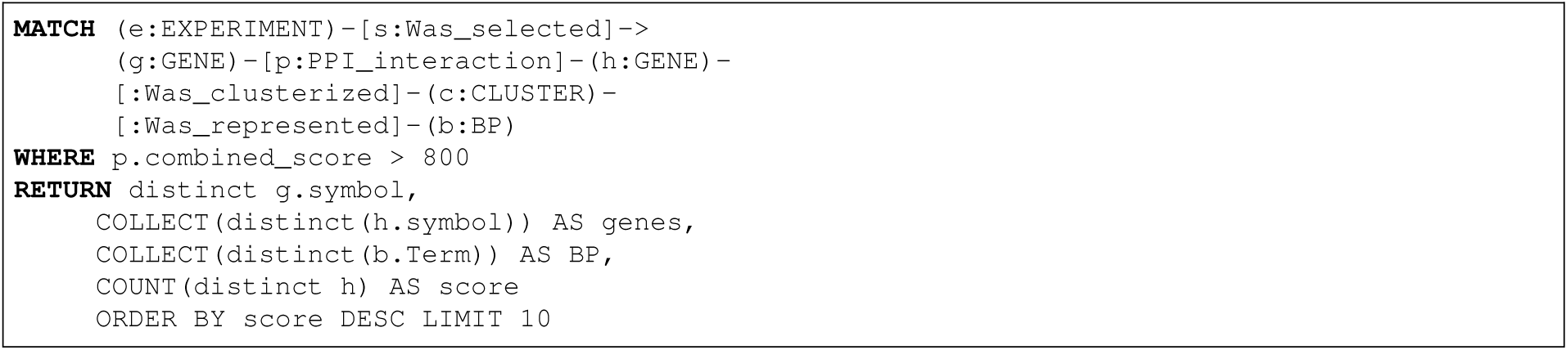
Which nodes of type TGENE were DE and present the highest number of connections associated to the protein interaction networks (PPI) according to a combined score value of > 0.80? Among these selected nodes, what are the clusters and associated biological processes?

Some common and important topological metrics in biological networks include: degree, distance, centrality, clustering coefficient. In this work, we use the degree metric *k_i_* of a node *n_i_*, defined as the number of edges that are adjacent (*a_ij_*) to this node, which is given by:

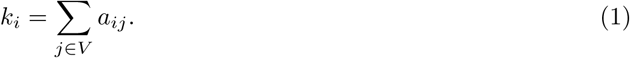

We use the Cypher query language to find the most connected *DE* genes in the network that establish known connections to the PPI network, having a high attribute value for the combined interaction score. For these genes we computed the co-expression cluster and, subsequently, the biological processes attributed to these clusters. One can observe that the query is expressed in a concise manner for answering a relatively complex topological question. The resulting DE genes are displayed in Table 2.

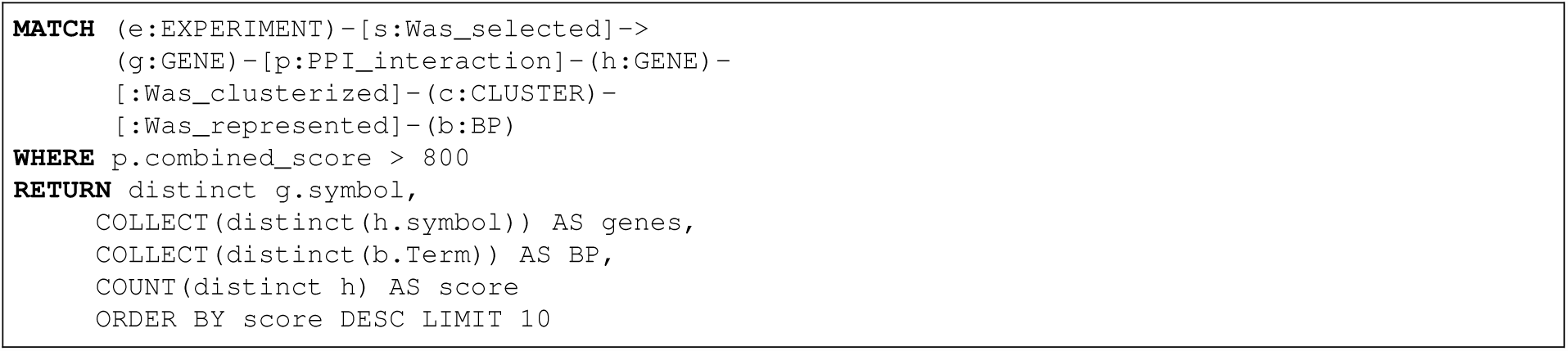

**Table 2:** Result showing the top 10 gene DE by PPI in experiment GSE28619. These genes are know as hubs and maybe are associated in important pathways in experimental context analyzed.

| Genes selected | cgn<sup>a</sup> | BP associated |
| --- | --- | --- |
| JUN | 89 | response to zinc ion; positive regulation of fibrinolysis; astrocyte activation; beta-alanine metabolic process; dibenzo-p-dioxin metabolic process; purine nucleobase catabolic process; positive regulation of gene expression; generation of precursor metabolites and energy; regulation of natural killer cell chemotaxis; ketone body biosynthetic process; |
| SRC | 83 |  |
| CDK1 | 80 |  |
| STAT3 | 75 |  |
| CREBBP | 72 |  |
| FOS | 62 |  |
| CDC42 | 58 |  |
| IGF1 | 58 |  |
| CCND1 | 55 |  |
| CDKN1A | 54 |  |

<sup>a</sup> number of connected genes

One of the main advantages of using the data model adopted in GeNNet is the availability of data and information that can be easily done without changing the data model. New nodes may add information such as metadata of samples (e.g. information on a patient’s eating habits) or new edges may add new relationships (e.g. genes co-expressed in different methods used) or even both (e.g. addition of a database on microRNA interactions connected to existing genes in the database). In the example below, we add a HUB-like node from the result obtained in query 3. Through the CREATE clause, after obtaining the selected genes, a new node and edges were created (Figure 7. This queries demonstrates the flexibility of the database in adding new information that can be generated through existing data in GeNNet-DB.

**Figure 7:**
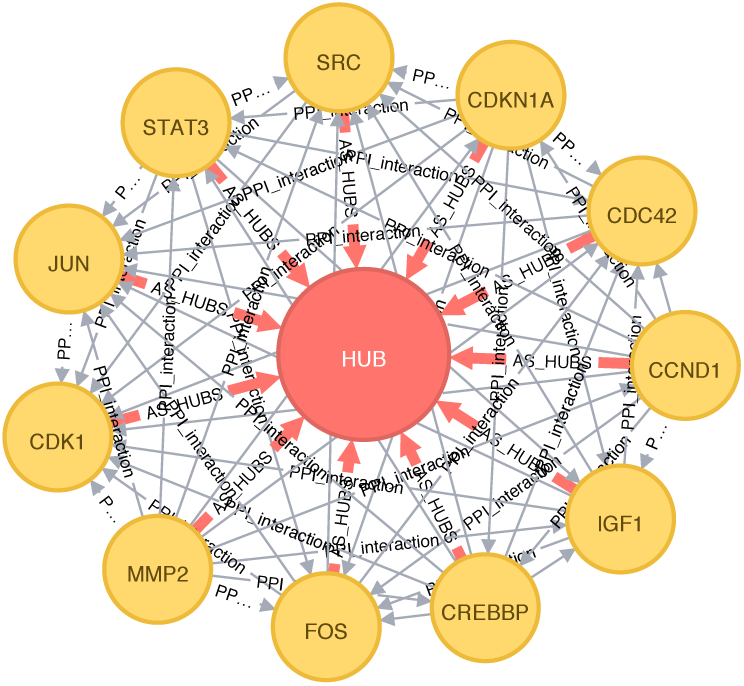
New nodes and edges added to the graph database. The genes that were highly connected according to query 3 were directed to the type node HUB.

**Query 4:**
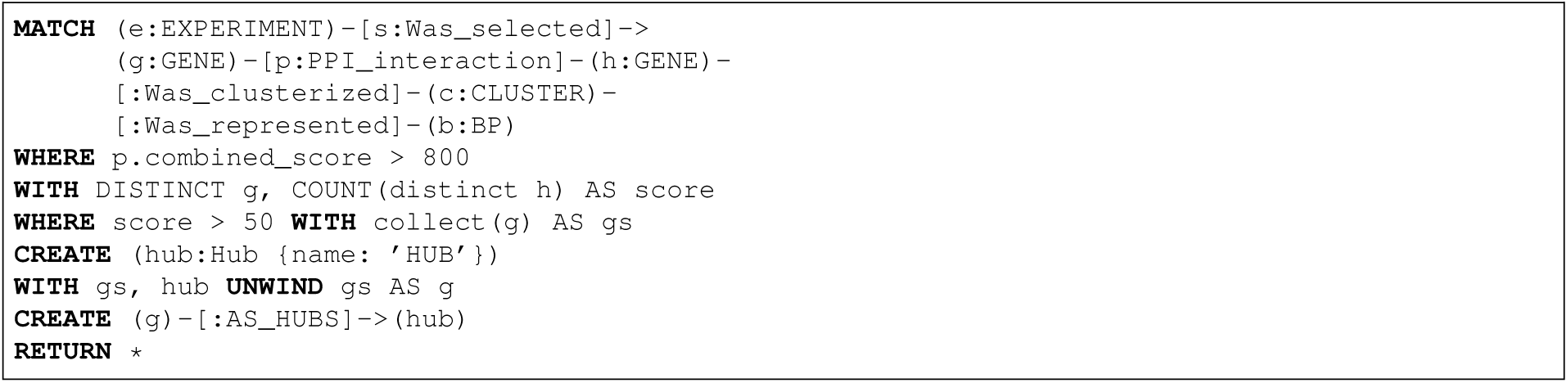
New node and edges inserted from the result of the previous query.

## Conclusion, Updates and Future Work

The platform presented in this work is the first one to integrate the analytical process of transcriptome data (currently only available for microarray essays) with graph databases. The results allow for testing previous hypothesis about the experiment as well as exploring new ones through the interactive graph database environment. It enables the analysis of different data coming from Affymetrix platforms on humans, rhesus, mice and rat.

The GeNNet will be periodically updated and we intend to extend the modules including analyses of RNA-seq and miRNA. We will incorporate additional experimental designs for DE and improve the execution time of the analyses. Due to the free access to GeNNet we rely on the feedback of the community for improving the tool. The distribution of the platform in a software container allows not only for executing it on a local machine but also to easily deploying it on a server and making it available on the Web.

## Acknowledgments

The authors thank all people’s contribution on this work.

## List of abbreviations

BP: Biological Process;
DE: Differential expression;
DDG: Data Derivation Graph;
EPEL: Extra Packages for Entrerprise Linux;
FDR: False Discovery Rate;
GeNNet-DB: database of the GeNNet
GeNNet-Web: web interface of the GeNNet;
GeNNet-Wf: workflow of the GeNNet;
GEO: Gene Expression Omnibus;
NoSQL: Not only SQL.

## Declarations

### Ethics approval and consent to participate

Not applicable.

### Consent for publication

Not applicable.

### Availability of data and materials

GeNNet’s source code is available at https://github.com/raquele/GeNNet. A software container that allows for easily executing GeNNet can be retrieved with the command `docker pull quelopes/gennet`.

### Authors’ contributions

RLC conceived idea, the software architecture, designed the graph query language and writing the manuscript. The LG worked in the software container implementation, the GeNNet architecture proposal, and on writing the manuscript. MRA and FP provided critical comments and improved upon the manuscript and application. All authors read and approved the final version of then manuscript.

## Footnotes

1 https://shiny.rstudio.com/

2 Available at: https://cran.r-project.org/web/packages/RNeo4j

3 https://www.docker.com

## Figures, tables additional files

### GeNNet tutorial

GeNNet tutorial is available at https://github.com/raquele/GeNNet.

### Some datasets using to test the database

**Table 3:**
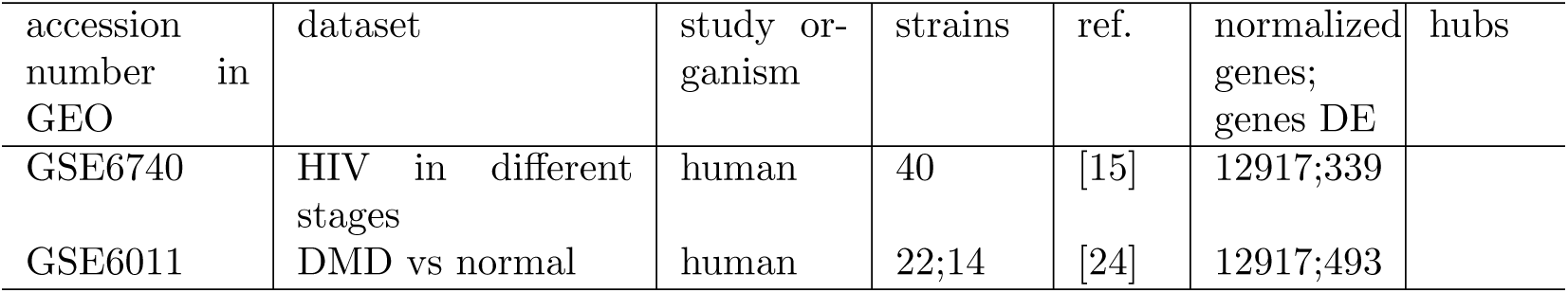
Some examples of gene expression experiments published in literature used in our platform.

| accession<br>number in<br>GEO | dataset | study or-<br>ganism | strains | ref. | normalized<br>genes;<br>genes DE | hubs |
| --- | --- | --- | --- | --- | --- | --- |
| GSE6740 | HIV in different<br>stages | human | 40 | [15] | 12917;339 |  |
| GSE6011 | DMD vs normal | human | 22;14 | [24] | 12917;493 |  |

